# Relative reward memories retain multidimensional information in bumblebees

**DOI:** 10.64898/2026.10.26.761910

**Authors:** Wenhao Zhang, Ke Wen, Tianheng Zhang, Yingwei Huang, Zhongmin Song, Yonghe Zhou, Cwyn Solvi, Fei Peng

**Affiliations:** State Key Laboratory of Multi-organ Injury Prevention and Treatment, Key Laboratory of Mental Health of the Ministry of Education, Guangdong-Hong Kong-Macao Greater Bay Area Center for Brain Science and Brain-Inspired Intelligence, Guangdong-Hong Kong Joint Laboratory for Psychiatric Disorders, Guangdong Provincial Key Laboratory of Psychiatric Disorders, Guangdong Basic Research Center of Excellence for Integrated Traditional and Western Medicine for Qingzhi Diseases, Guangdong Provincial Key Laboratory of Tropical Disease Research, Department of Psychology, School of Public Health, Southern Medical University, Guangzhou, China; Department of Psychiatry, Zhujiang Hospital, Southern Medical University, Guangzhou, China; Department of Neurobiology, School of Basic Medical Sciences, Southern Medical University, Guangzhou, China; Institute of Artificial Intelligence, School of Future Technology, Shanghai University, Shanghai, China

## Abstract

Previously, we showed that bumblebees encode reward value relative to local alternatives and, when comparing options learned in separate contexts, retrieve only their ordinal rank rather than absolute reward magnitude (Solvi et al., 2022). However, that study, like related work across species, varied reward along a single dimension. Natural rewards such as nectar are multidimensional, yet how such rewards are represented in relative memory remains unknown. We tested this by training bumblebees in separate contexts with rewards varying in two intrinsic dimensions, sucrose concentration and NaCl content, and then comparing options never encountered together. When only sucrose concentration varied, bees showed no preference between options that had each been locally better, despite differing in absolute concentration. With multidimensional rewards, however, this equivalence no longer held: bees preferred an option that had been locally better on both sucrose and NaCl over one better on only sucrose or only NaCl. A two-dimensional advantage also produced a stronger preference than a one-dimensional advantage 48 h after training. Thus, the ordinal rank we previously identified captures only part of the relative reward representation: with multidimensional rewards, bumblebees also retain information about the dimensions on which an option had been locally better.

## Introduction

What do animals remember about the value of options outside the contexts in which they were learned? Across species, remembered option values are shaped by the alternatives available during learning, such that later choices between options learned in different contexts can deviate from their absolute reward values (Bavard et al., 2021, 2018; Ferguson et al., 2026; Klein et al., 2017; Palminteri et al., 2015; Pompilio and Kacelnik, 2010; Solvi et al., 2022). In our previous study, bumblebees showed a particularly clear form of such relative reward memory. Bees learned that one option was better than another in one context (A > B) and that a different option was better than another in a second context (C > D). When options from the two contexts were later compared (A vs. C) for the first time, bees treated options with the same local rank as equivalent (A = C) despite differences in absolute reward magnitude (Solvi et al., 2022).

However, like related work across species, that study varied reward along a single dimension. Natural rewards, by contrast, are multidimensional, with multiple properties simultaneously contributing to reward value. How such multidimensional rewards are represented in relative memory remains unknown across species, including in humans (Ferguson et al., 2026; Palminteri and Lebreton, 2021).

To address this question, we extended our previous paradigm to include two intrinsic reward dimensions, sucrose concentration and NaCl content. We asked whether options with the same local rank would remain equivalent when one had been locally better on two dimensions and the other on only one dimension. If not, this would indicate that relative memory preserves information beyond an option’s overall local rank, including the reward dimensions on which it had been locally better.

## Results

### A one-dimensional baseline for relative reward memory

We first asked whether the ordinal rank memory reported previously (Solvi et al., 2022) would be reproduced under the present experimental conditions, thereby establishing a one-dimensional baseline for the multidimensional reward experiments. In Experiment 1, bees (N = 20) foraged individually from artificial flowers in a flight arena (see supplementary materials for full methodological details). Each bee learned two choice contexts in counterbalanced order, with each option identified by a distinct visual pattern (Figure 1A, B). In context AB, A provided 45% (w/w) sucrose and B 30%, whereas in context CD, C provided 30% sucrose and D 20%. Thus, A and C shared the same local ordinal rank – each was better than its local alternative (A > B and C > D) – despite differing in absolute sucrose concentration. After training in both contexts, each bee encountered A and C together for the first time in an unrewarded transfer test (Figure 1C).

**Figure 1.**
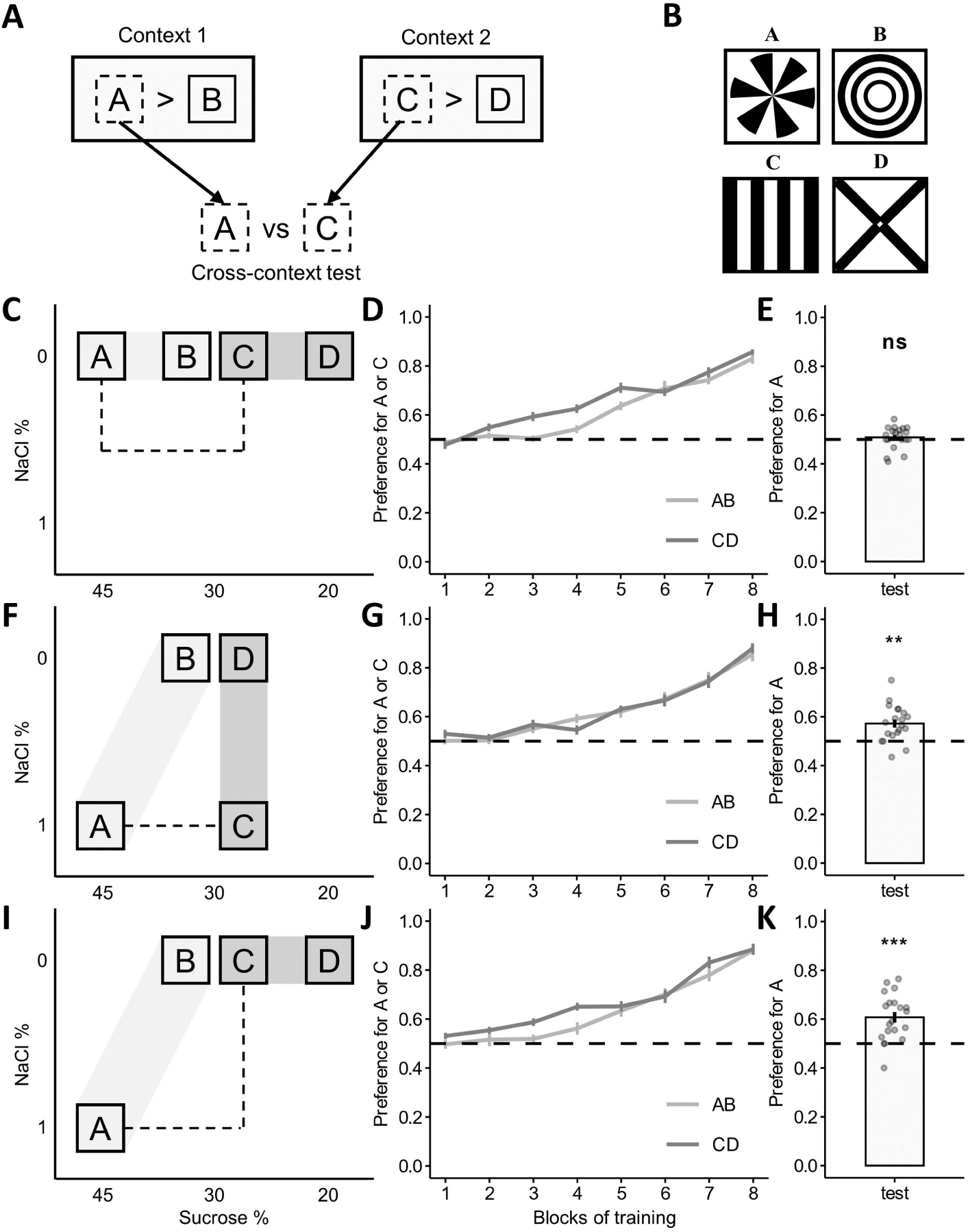
Reward dimensions supporting local advantage determine cross-context choice. (A) General training and transfer procedure used in Experiments 1–3. Each bee learned that A was better than B in one context and C was better than D in another. The two locally better options, A and C, were then presented together for the first time in an unrewarded cross-context transfer test. (B) Achromatic visual patterns identifying options A–D. Pattern assignment was fixed across bees and experiments. (C) Design of Experiment 1. A was better than B in sucrose concentration (45 versus 30%), and C was better than D in sucrose concentration (30 versus 20%). (D) Bees learned to prefer the locally better option in both contexts. (E) Bees showed no detectable preference between A and C in the unrewarded transfer test. (F) Design of Experiment 2. A was better than B in both sucrose concentration and NaCl content, whereas C was better than D in NaCl content alone. (G) Bees learned both discriminations. (H) Bees preferred A over C in the transfer test. (I) Design of Experiment 3. A was better than B in both sucrose concentration and NaCl content, whereas C was better than D in sucrose concentration alone. (J) Bees learned both discriminations. (K) Bees again preferred A over C in the transfer test. In (C), (F), and (I), shaded bands connect options trained together and dashed lines connect options presented in the transfer test. Training blocks ended after 10 drinking visits to the locally better option. Lines and bars show means, error bars show SEM, and circles represent individual bees. Horizontal dashed lines indicate chance. N = 20 bees per experiment. Transfer preferences were tested using generalized linear mixed models. ns, P > 0.05; **P < 0.01; ***P < 0.001.

Training in each context continued until each bee had completed 80 drinking visits to the locally better option (A or C), thereby equating rewarded experience with the two later transfer options. To visualize learning, choices within each context were divided into eight blocks, each ending after 10 drinking visits to A or C (Figure 1D).

Preference was calculated as the proportion of choices within each block directed toward the locally better option against its alternative (B or D). Preference increased during training in both contexts. By the final training block, bees preferred A over B (generalized linear mixed model [GLMM]: Z = 9.093, P < 2e-16) and C over D (GLMM: Z = 9.552, P < 2e-16), confirming that both local discriminations had been learned. Preference strength did not differ detectably between contexts (LRT: χ² = 1.093, P = 0.296) and was not affected by which context was trained first (LRT: χ² = 0.705, P = 0.401). In the subsequent transfer test, bees showed no detectable preference between A and C (GLMM: Z = 0.525, P = 0.600; Figure 1E). Thus, Experiment 1 reproduced the previous ordinal memory result: despite differing in absolute sucrose concentration, options sharing the same local rank were treated as equivalent.

### Multidimensional rewards break ordinal equivalence

Having established the equivalence between locally better options when sucrose was the only varying reward dimension, we introduced NaCl as a second reward dimension. Sodium occurs naturally in floral nectar (Nicolson, 2022), can influence bees’ feeding preference (Sousa et al., 2022), and, when enriched in nectar, increases pollinator visitation rate and diversity (Finkelstein et al., 2022; VanValkenburg et al., 2024), supporting NaCl as a biologically relevant reward dimension. In Experiment 2 (N = 20; Figure 1F), both options later compared in the transfer test had contained 1% (w/w) NaCl during training. A (45% sucrose + 1% NaCl) was trained against B (30% sucrose), whereas C (30% sucrose + 1% NaCl) was trained against D (30% sucrose). Thus, A was locally better than B in both sucrose concentration and NaCl content, whereas C was locally better than D in NaCl content alone. By the final block, bees preferred A over B (GLMM: Z = 6.950, P = 3.64e-12) and C over D (GLMM: Z = 9.690, P < 2e-16), confirming that 1% NaCl could support learned preference when sucrose concentration was held constant. Final-block preference did not differ detectably between contexts (LRT: χ² = 0.934, P = 0.334) or with contextual training order (LRT: χ² = 0.302, P = 0.583; Figure 1G).

Nevertheless, bees preferred A over C in the unrewarded transfer test (GLMM: N = 20, Z = 2.795, P = 0.005; Fig. 1H). This preference could not be attributed simply to NaCl presence, because A and C had both contained 1% NaCl and been rewarded equally often. Nor could absolute sucrose concentration alone explain it: bees showed no detectable preference for the same 45%-versus-30% comparison in Experiment 1. A further possibility was that NaCl did not act as a separate reward dimension, but instead contributed to the perceived value of sucrose, functionally similar to increasing sucrose concentration. If so, A and C would both have been encoded simply as locally better options, between which bees should show no preference, as observed in Experiment 1 and in our previous study (Solvi et al., 2022). Their later preference for A therefore argues against this possibility, and suggests that NaCl contributed reward information distinguishable from sucrose.

Experiment 2 showed that an option locally better on two dimensions was preferred over one locally better on NaCl alone. We next asked whether the same pattern would hold when the one-dimensional option was instead better on sucrose. In Experiment 3 (N = 20; Figure 1I), A (45% sucrose + 1% NaCl) was trained against B (30% sucrose), whereas C (30% sucrose) was trained against D (20% sucrose). Thus, both A and C were locally better on sucrose, but only A was additionally better on NaCl.

Bees reached comparable preference for the locally better option (A or C) by the final training block (LRT: χ² = 1e-04, P = 0.993), with no detectable effect of training order (LRT: χ² = 0.219, P = 0.640; Figure 1J). Nevertheless, bees again preferred A over C in the subsequent transfer test (GLMM: N = 20, Z = 4.229, P = 2.35e-05; Figure 1K). Moreover, the strength of the preference for A did not differ detectably between Experiments 2 and 3 (LRT: χ² = 1.260, P = 0.262), providing no evidence that the effect depended on whether the one-dimension option was better on sucrose or NaCl. Together, Experiments 2 and 3 show that an option that had been locally better on two dimensions was preferred over one that had been locally better on one dimension, whether that dimension was NaCl or sucrose.

### A multidimensional advantage strengthens long-term preference

Having found that an advantage on two reward dimensions influenced cross-context choice, we asked whether it also affected long-term preference (Figure 2A). In Experiment 4, one group of bees (N = 12) learned that A was better than B in both sucrose concentration and NaCl content (A: 45% sucrose + 1% NaCl; B: 30% sucrose), whereas another group (N = 12) learned that A was better in sucrose concentration alone (A: 45% sucrose; B: 30% sucrose; Figure 2B). Training ended after 60 rewarded visits to A, and bees received an unrewarded test 48 hours later. Final-block preference for A did not differ detectably between groups (LRT: χ² = 1.277, P = 0.258; Figure 2C). After 48 hours, however, bees trained with the two-dimensional advantage preferred A more strongly than those trained with the sucrose-only advantage (LRT: χ² = 5.643, P = 0.018; Figure 2D), suggesting that a multidimensional reward advantage can support stronger long-term preference.

**Figure 2.**
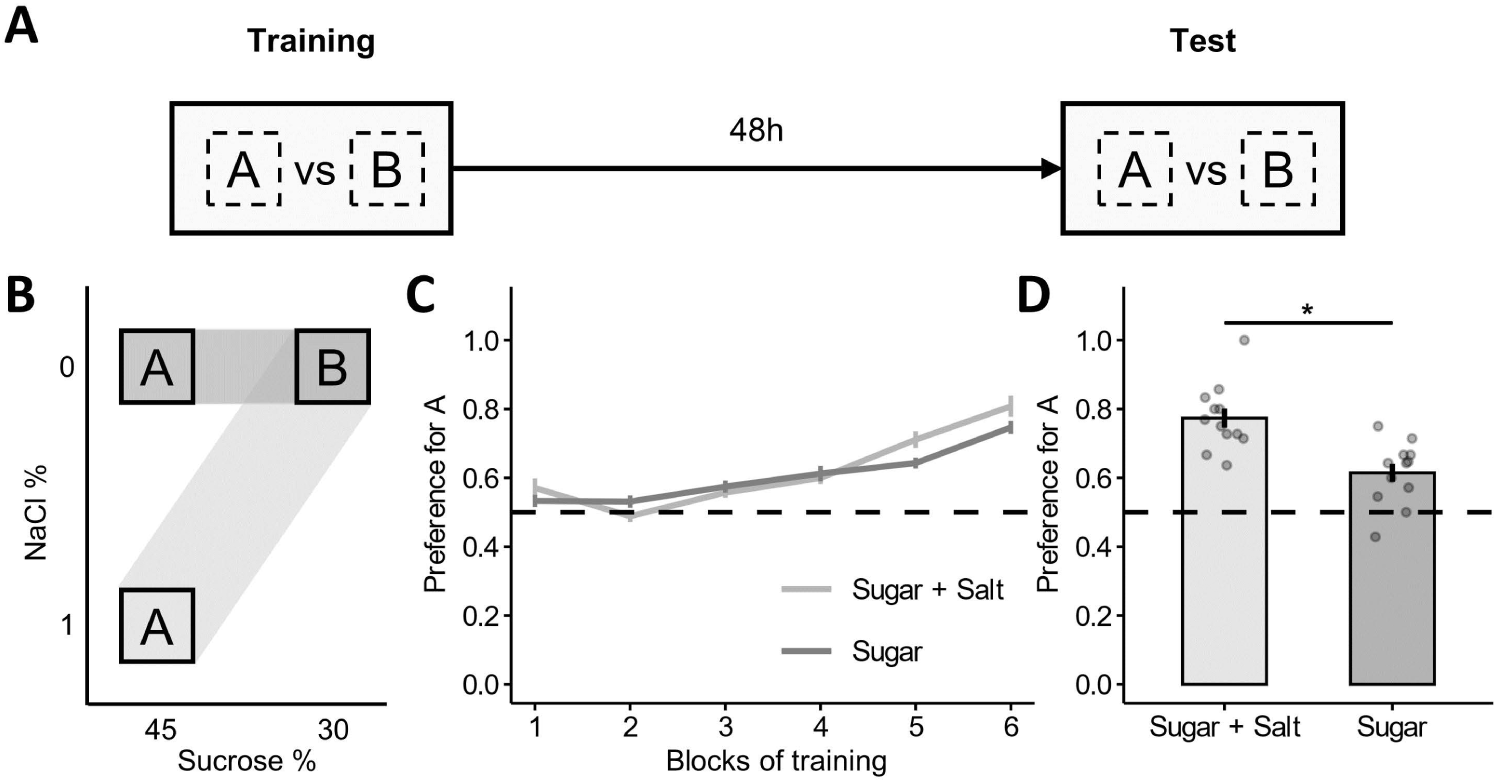
Multidimensional reward strengthens long-term preference. (A) Procedure for Experiment 4. Bees were trained to discriminate A from B and tested with the same options without reward 48 hours later. (B) Experimental design (N = 12 per group). In the sugar-and-salt group, A was better than B in both sucrose concentration and NaCl content; in the sugar-only group, A was better than B in sucrose concentration alone. Shaded bands connect options trained together. (C) The two groups expressed comparable preferences for A during training. (D) After 48 hours, bees trained with the two-dimensional advantage showed a stronger preference for A than the one-dimensional group. Lines and bars show means, error bars show SEM, and circles represent individual bees. Horizontal dashed lines indicate chance. *P < 0.05.

## Discussion

Our previously study showed that when only sucrose concentration varied, bumblebees showed no detectable preference between locally better options despite differences in absolute concentration (Solvi et al., 2022). Here we first reproduced the equivalence between locally better options when sucrose concentration was the only varying reward dimension, and then showed that this equivalence broke down when rewards were multidimensional: an option that had been locally better on both sucrose concentration and NaCl content was preferred in a later cross-context comparison over one that had been better on only one dimension, regardless of whether that dimension was sucrose or NaCl. Thus, the ordinal rank revealed by one-dimensional tasks captures only part of the relative reward representation. Relative memory can lose information about how much better an option was while retaining information about the dimensions on which it had been locally better.

Our results also raise a question about the order of operations: are different reward dimensions combined before or after they are evaluated relative to context? A common view of multidimensional choice is that the dimensions of each option are first combined into a single overall value, which is then compared with the value of other options (Lim et al., 2013). An alternative is that corresponding dimensions are first compared across options, and their contributions are combined only afterward (Amasino et al., 2019; Hunt et al., 2014; Perkins et al., 2024). Our results extend this distinction from choices between simultaneously available options to memories of outcomes learned in separate contexts. If sucrose and NaCl were first combined into a single value, and that value was then reduced to the same context-relative representation observed in the one-dimensional baseline shown in Experiment 1 and previous work (Solvi et al., 2022), A and C should have remained equivalent in Experiments 2 and 3. Instead, bees preferred A over C in both experiments. This pattern is consistent with that sucrose and NaCl being evaluated separately relative to their local alternatives before their contributions were combined in the subsequent cross-context comparison.

One candidate process that could generate this pattern is Decision by Sampling. Rather than representing how much better one outcome is than another, Decision by Sampling makes a series of simple ordinal comparisons – whether an option is better or worse than another – and subjective value emerges from how often an option wins these comparisons (Stewart et al., 2006; Vlaev et al., 2011). Applied separately across reward dimensions, such comparisons could establish whether an option had been locally better in sucrose or NaCl, with the outcomes of these comparisons later combined when options from different contexts were encountered together.

Multidimensional extensions of Decision by Sampling formalize this possibility (Noguchi and Stewart, 2018) and related rank-based models capture context-dependent transfer choices in human reinforcement learning (Hayes and Wedell, 2023).

A reward representation that preserves multidimensional structure may be particularly useful in fluctuating, multidimensional environments. Natural nectar, for example, varies in sugar concentration, salt content, and other constituents (Nicolson, 2022), creating an inherently multidimensional reward. Together with evidence that nectar constituents can strengthen pollinator memory and floral fidelity (Hemingway et al., 2024), our 48-hour result raises the possibility that reward complexity can strengthen longer-term flower preferences. This ecological possibility also connects to a broader computational principle. In feature-based learning and multi-objective reinforcement learning, retaining information about different outcome dimensions can support flexible choice when goals or priorities change, because their contributions can be recombined or reweighted rather than requiring option values to be learned anew (Farashahi et al., 2017; Hayes et al., 2022; Roijers et al., 2013; Shiravand et al., 2026). The structure retained by context-dependent reward memory could support similar flexibility.

This broader perspective may also shed light on a longstanding functional puzzle (Palminteri and Lebreton, 2021): why should relative encoding be so widespread when it can distort choices between options learned in different contexts? Existing accounts emphasize the benefits of adapting value representations to local reward ranges or distributions, which can improve learning efficiency at the cost of accurate cross-context comparison (Bavard et al., 2018; Bavard and Palminteri, 2023; Bhui and Gershman, 2018; Glimcher, 2022; Hunter and Daw, 2021). Our results suggest that one-dimensional tasks may overstate the cost of relative encoding: absolute reward magnitude can be lost while multidimensional structure is preserved. Viewed in this light, what appears irrational in a one-dimensional task may therefore reflect a representation adapted to richer, multidimensional rewards (Fawcett et al., 2014; Summerfield and Parpart, 2021).

## Materials and Methods

### Animals

Colonies of bumblebees (Bombus terrestris) were obtained from a commercial supplier (Biobest Belgium N.V., Westerlo, Belgium) and housed in wooden nest boxes (28 cm × 16 cm × 11 cm). Although there are no current requirements regarding insect care and use in research, we refined our methods to minimize any stress on the bees and reduced the number of bees and colonies used as much as possible. In addition, the behavioral tests were non-invasive, and the substances used (sucrose, NaCl, and water) occur naturally in the foraging environment of bumblebees. Bumble bees were cared for daily by trained and competent staff, including routine welfare and provision of correct and adequate food during the experimental period. Outside of experimental work, bees had ad libitum access to 20% sucrose solution (w/w; all concentrations below are expressed as w/w) and were also supplied with approximately 3 g pollen every day.

### Experimental setup

Bees were trained and tested individually in a flight arena (40 cm × 59 cm × 41 cm) connected to the nest box by a transparent acrylic tunnel (30 cm × 4 cm × 4 cm) fitted with sliding gates, allowing individual foragers to enter the arena.

The top of the arena was covered by a sheet of transparent UV-transmitting acrylic. A USB industrial vision camera (ZWAK 5600, Zhongweiaoke, China) was installed in the center of the acrylic ceiling to record all experiments from a top-down view.

Each artificial flower (hereafter, “flower”) consisted of an acrylic chip (25 mm x 25 mm x 5 mm) mounted on an opaque glass cylinder (4cm high). During pre-training, all flowers had transparent chips. During training and testing, the chips were covered with achromatic printed patterns (30 mm x 30 mm). The four visual patterns shown in Fig. 1 were consistently assigned to options A, B, C, and D, and were not counterbalanced. Preliminary observations indicated no detectable spontaneous preference among the patterns and choices were at chance level at the beginning of training across experiments.

Reward contingencies differed among experiments. In Experiment 1, bees (N = 20) were individually trained with two one-dimensional reward contrasts: in context AB, A contained 45% sucrose and B contained 30% sucrose; in context CD, C contained 30% sucrose and D contained 20% sucrose. In Experiment 2 (N = 20), A contained 45% sucrose supplemented with 1% NaCl and B contained 30% sucrose, whereas C contained 30% sucrose supplemented with 1% NaCl and D contained 30% sucrose alone. In Experiment 3 (N = 20), A contained 45% sucrose supplemented with 1% NaCl and B contained 30% sucrose, whereas C contained 30% sucrose and D contained 20% sucrose. In Experiment 4, the two-dimensional group (N = 12) was trained with A containing 45% sucrose supplemented with 1% NaCl and B containing 30% sucrose, whereas the one-dimensional control group (N = 12) was trained with A containing 45% sucrose and B containing 30% sucrose.

### Experiment procedure

Bees were individually pre-trained to forage from eight transparent flowers. All flowers were filled with 10 µL 20% sucrose solution. Depleted rewards were replenished after the bee landed on a subsequent flower. After each foraging bout, flowers were cleaned with 75% ethanol and repositioned pseudo-randomly. Bees that completed three consecutive foraging bouts were classified as foragers and were marked with number tags (Opalithplättchen, Warnholz& Bienenvoigt, Ellerau, Germany) attached to their thorax with a minimal amount of superglue. Only identified foragers proceeded to formal training.

### Training and testing

Experiments 1-3 followed the same contextual training and out-of-context transfer test procedure. Each bee completed two training phases, one in context AB and one in context CD. In each phase, bees foraged from an array of eight flowers, four displaying each of the two visual patterns used in that context. One option provided the locally superior reward and the other provided the locally inferior reward. A choice was recorded when a bee landed on a flower and extended its proboscis to contact the solution. During all training phases, depleted rewards were replenished during a bout, and flowers were cleaned with 75% ethanol and repositioned pseudorandomly between bouts. Training in each context ended after the bee had made 80 choices of the locally superior pattern. The two training phases were separated by one hour, and their order was counterbalanced across bees. During training, each bee learned to prefer flower A over B in one context, and C over D in another. One hour after completing the second training phase, each bee received a 2-min unrewarded transfer test between A and C. During the test, each flower contained 10 µL of distilled water.

In Experiment 4, bees similarly completed a single training phase with options A and B. Training ended after 60 choices of A had been made. Bees received a 2-min unrewarded test with the same two options 48 h after training, with 10 µL of distilled water provided on each flower.

### Statistical analysis

All statistical analyses were conducted in R (v. 4.4.1). Generalized linear mixed models (GLMMs) were fitted using glmmTMB (v.1.1.10), with a binomial error distribution and logit link function. Responses were specified as choices for the focal option versus its alternative, with the total number of choices providing the binomial denominator. Training choices were grouped into blocks of 10 visits to the focal option for visualization (eight blocks in Experiments 1–3 and six in Experiment 4). In Experiments 1–3, training context and context order were tested using final-block choices, whereas transfer choices between A and C were compared with chance using intercept-only models; experiment identity was included as a fixed effect when comparing transfer preferences between Experiments 2 and 3. In Experiment 4, group was included as a fixed effect in separate comparisons of final-block and 48-hour test choices. Bee identity was included as a random effect. Fixed effects were evaluated using likelihood-ratio tests (LRTs) comparing nested models, whereas deviations from chance were assessed using Wald z tests of the intercept.

